# Overexpression of CD47 is associated with brain overgrowth in 16p11.2 deletion syndrome

**DOI:** 10.1101/808022

**Authors:** Jingling Li, Thomas Brickler, Allison Banuelos, Kristopher Marjon, Jing Bian, Cyndhavi Narayanan, Irving L. Weissman, Sundari Chetty

## Abstract

One of the most common genetic linkages associated with neuropsychiatric disorders, such as autism spectrum disorder and schizophrenia, occurs at the 16p11.2 locus. Copy number variants (CNVs) of the 16p gene can manifest in opposing head sizes. 16p11.2 deletion carriers tend to have macrocephaly (or brain enlargement), while those with 16p11.2 duplication frequently have microcephaly. Increases in both gray and white matter volume have been observed in brain imaging studies in 16p11.2 deletion carriers with macrocephaly. Here, we use human induced pluripotent stem cells (hiPSCs) derived from controls and subjects with 16p11.2 deletion and 16p11.2 duplication to understand the underlying mechanisms regulating brain overgrowth. To model both gray and white matter, we differentiated patient-derived iPSCs into neural progenitor cells (NPCs) and oligodendrocyte progenitor cells (OPCs). In both NPCs and OPCs, we show that CD47 (a ‘don’t eat me’ signal) is overexpressed in the 16p11.2 deletion carriers contributing to reduced phagocytosis both *in vitro* and *in vivo*. Treatment of 16p11.2 deletion NPCs and OPCs with an anti-CD47 antibody to block CD47 restores phagocytosis to control levels. Furthermore, 16p11.2 deletion NPCs and OPCs upregulate cell surface expression of calreticulin (a pro-phagocytic ‘eat me’ signal) and its binding sites, indicating that these cells should be phagocytosed but fail to be eliminated due to elevations in CD47. While the CD47 pathway is commonly implicated in cancer progression, we document a novel role for CD47 in regulating brain overgrowth in psychiatric disorders and identify new targets for therapeutic intervention.

## Introduction

Autism spectrum disorder (ASD) is a complex neurodevelopmental disorder, characterized by deficits in social interaction and communication and frequently associated with restrictive, repetitive behaviors. ASD is the fastest growing developmental disability, affecting 1 in 59 children in the United States. Heterogeneity and access to relevant cell types have made it challenging to understand underlying mechanisms and identify effective therapies for ASD. 16p11.2 copy number variation (CNV) is one of the most common genetic linkages of ASD as well as other intellectual disabilities and developmental delays^1–7^.

A large head size (referred to as macrocephaly) is common in people who have 16p11.2 deletion syndrome. Head circumference is highly correlated with brain size early in development^8^. While macrocephaly refers to having enlarged head circumference (typically greater than the 97th percentile), microcephaly refers to having small head circumferences below the 3^rd^ percentile^9^. Meta-analyses studies have found that approximately 15-20% of individuals with ASD have macrocephaly^9,10^, with disproportionate enlargement in both gray and white matter volume^11,12^. Individuals with ASD and macrocephaly have more severe behavioral and cognitive problems--such as lower language ability and slower cognitive development into early childhood--and are less responsive to standard medical and therapeutic interventions, leading to very poor prognoses relative to individuals with ASD and normal head circumferences^13^. Increases in brain size often precede clinical symptoms^12–14^, suggesting that understanding the underlying mechanisms regulating brain overgrowth could provide a window of opportunity for intervention or mitigation of symptoms. It is important to use human-derived samples to better understand the disorder^15^, because 16p11.2 CNV in mouse models affects a different set and number of genes within the 16p locus and has the opposite phenotype relative to humans—enlarged brains in 16p11.2 duplication and microcephalic brains in the 16p11.2 deletion syndrome^16,17^.

Here, we focus on a clinically meaningful subtype of individuals with intellectual disability (IQ<70) or ASD associated with brain overgrowth in 16p11.2 deletion carriers. We use patient-derived human induced pluripotent stem cells (hiPSCs) to interrogate the underlying mechanisms contributing to gray and white matter enlargement. Specifically, we differentiate the iPSCs into neural progenitor cells (NPCs) and oligodendrocyte progenitor cells (OPCs) and investigate the hypothesis that brain enlargement in 16p11.2 deletion carriers may be due to improper cellular elimination. Under normal conditions, classical ‘eat me’ and ‘don’t eat me’ signaling mechanisms associated with phagocytosis maintain cellular homeostasis across diverse tissue types^18,19^. We investigated whether these signaling mechanisms may be deregulated in conditions associated with brain overgrowth. We show that CD47 (a ‘don’t eat me’ signal) is overexpressed in NPCs and OPCs derived from 16p11.2 deletion carriers leading to reduced phagocytosis by macrophages and microglia. Furthermore, treatment with a CD47 blocking antibody can restore phagocytosis of 16p11.2 deletion NPCs and OPCs to control levels. Importantly, the 16p11.2 deletion NPCs and OPCs have increased cell surface expression of calreticulin (a pro-phagocytic ‘eat me’ signal), indicating that these cells should be eliminated but are not due to high levels of CD47^20^. We identify a novel role for CD47 in regulating brain size and highlight the importance of blocking CD47 to promote clearance of unhealthy NPCs and OPCs in 16p11.2 deletion with brain overgrowth.

To our knowledge, this is the first study to uncover a role for CD47 in neurodevelopmental disorders, such as ASD. A recent genome-wide association study (GWAS) has identified CD47 as a significant risk loci for bipolar disorder^21^, providing support that CD47 may play critical roles in brain regulation and function. Consistent with recent work on the role of CD47 in regulating synaptic pruning in brain development^22^ and social behavior in mouse models^23^, our study further highlights that the balance between ‘eat me’ and ‘don’t eat me’ signals may be more broadly playing critical roles during early neurodevelopment.

## Results

### Upregulation of CD47 in 16p11.2 deletion NPCs suppresses phagocytosis

We obtained human iPSCs from the Simons Foundation Autism Research Initiative (SFARI) from individuals who are carriers of the 16p11.2 CNV with complete clinical and phenotypic data. To better understand and elucidate mechanisms underlying brain overgrowth, we focused our study on 16p11.2 deletion carriers with defined macrocephaly (mean head circumference of 97± 2 percentile; three 16p11.2 del subjects had head circumferences in the 99th percentile, and one 16p11.2 del subject had a head circumference in the 91st percentile) (Figure 1A, Supplementary Table 1). All four of these 16p11.2 del subjects have been diagnosed with intellectual disability (IQ<70) or ASD (Supplementary Table 1) as well as phonological and/or language disorders. Two 16p11.2 deletion carriers with normal head circumferences without macrocephaly (Supplementary Table 1) were also used as comparison to identify whether our findings are specific to 16p11.2 deletion or 16p11.2 deletion with brain overgrowth. 16p11.2 duplication carriers in our study have head circumference percentiles within the normal range of 42.5±30.5 percentile with no diagnosis of macro- or microcephaly (Figure 1A, Supplementary Table 1). iPSCs from control subjects were obtained through the NIH registry with no genetic abnormalities.

**Figure 1.**
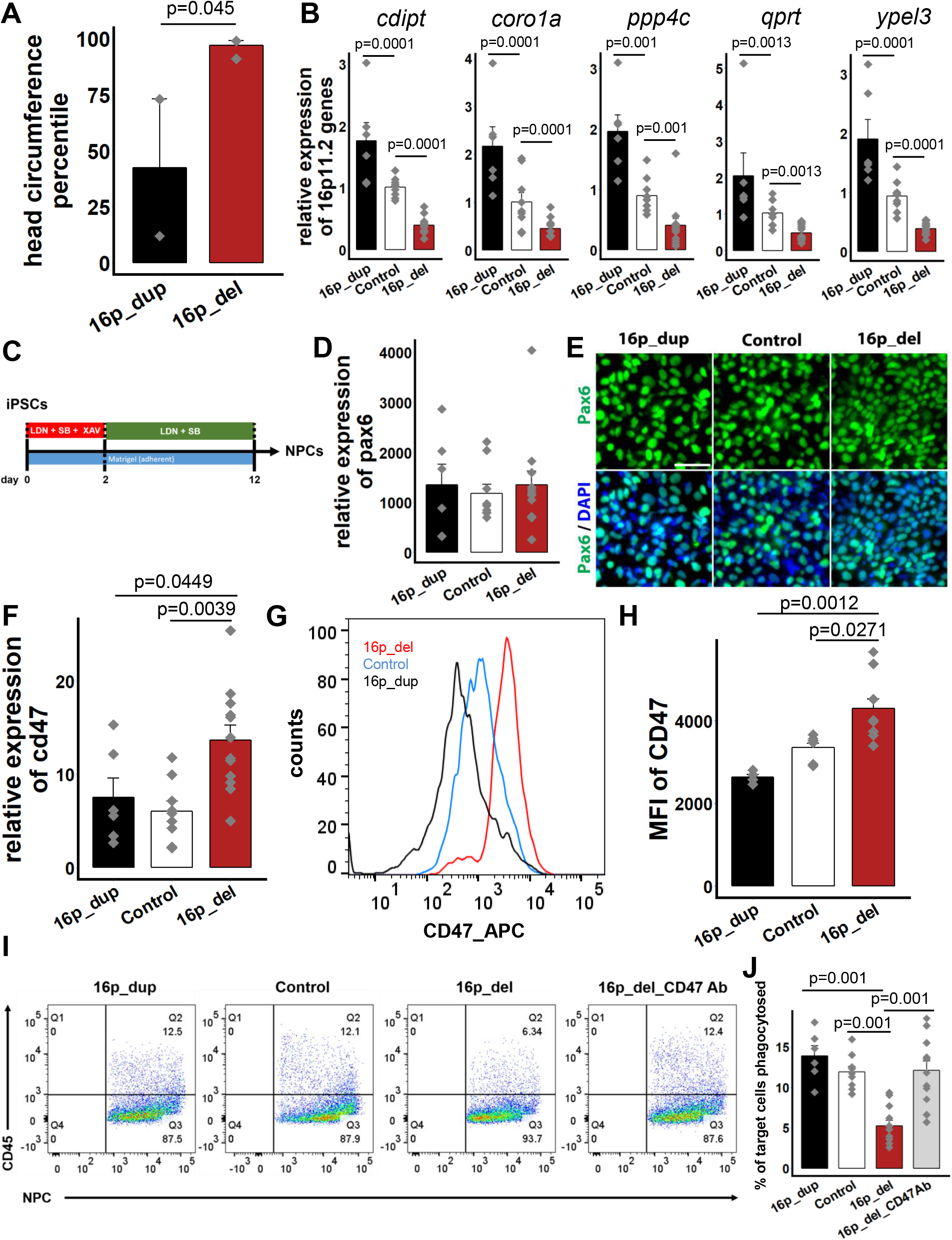
CD47 is overexpressed in 16p11.2 deletion NPCs. **A.** Head circumference measurements of 16p11.2 deletion (16p_del) and 16p11.2 duplication (16p_dup) carriers. 16p_dup, n=2; 16p_del, n=4. P value was determined by two-tailed equal variance student’s *t*- test. **B.** Quantitative RT-PCR shows an increase in expression of genes within the 16p11.2 locus in the 16p_dup iPSC lines and a reduction in 16p_del iPSC lines relative to control iPSC lines. Data represent fold change relative to undifferentiated control human iPSCs. n = 3 biological replicates per cell line (16p_dup, n=2; control, n=3; 16p_del, n=4 cell lines). P values were determined by one-way ANOVA followed by *post-hoc* Tukey HSD Test. **C.** Schematic figure showing directed differentiation of iPSCs into neural progenitor cells (NPCs). **D.** mRNA expression levels for *pax6* in 16p_dup, 16p_del, and control lines after 12 days of directed differentiation. Data represent fold change relative to undifferentiated control human iPSCs. n = 3 biological replicates per cell line (16p_dup, n=2; control, n=3; 16p_del, n=4 cell lines). Mean ± SE subjected to a one-way ANOVA followed by *post-hoc* Tukey HSD Test. **E.** Immunostaining for the NPC marker Pax6 in NPCs differentiated from 16p_dup, 16p_del, and control lines. Scale bar: 50 μm. **F.** Quantitative RT-PCR of *cd47* mRNA expression in 16p_del NPCs compared to control and 16p_dup NPCs. Data represent fold change relative to undifferentiated control human iPSCs. n = 3 biological replicates per cell line (16p_dup, n=2; control, n=3; 16p_del, n=4 cell lines). **G.** Representative flow cytometry histograms showing the mean fluorescence intensities (MFI) of CD47 in NPCs differentiated from 16p_dup, 16p_del, and control iPSCs. **H.** Quantification of CD47 MFI of the CD47 protein levels in 16p_del NPCs compared to control and 16p_dup NPCs. n = 2 biological replicates per cell line (16p_dup, n=2; control, n=3; 16p_del, n=4 cell lines). **I.** Representative flow cytometry phagocytosis plots showing rates of engulfment of NPCs when co-cultured with human derived macrophages. Phagocytosis assays were conducted using CellTrace Far Red-labeled 12 day differentiated NPCs and human macrophages. A CD45 antibody conjugated with PE was used to specifically label human macrophages. NPCs differentiated from 16p_del lines were also pretreated with or without a CD47 blocking antibody (CD47 Ab) prior to phagocytosis assessment. **J.** The percentages of phagocytosed NPCs in 16p_del lines compared to control and 16p_dup lines. Rates of phagocytosis of NPCs in the 16p_del lines following anti-CD47 treatment with a CD47 blocking antibody (16p_del_CD47Ab) are also shown. n=3 biological replicates per cell line (16p_dup, n=2; control, n=3; 16p_del, n=4 cell lines). All data are mean ± SE. P values were determined by one-way ANOVA followed by *post-hoc* Tukey HSD Test (**F**, **H**, **J**).

To confirm CNV status of the iPSCs, we first assessed mRNA levels of a subset of genes spanning the 16p11.2 locus by quantitative RT-PCR (qRT-PCR). Expression levels of many genes within this locus were significantly upregulated in the 16p11.2 duplication (16p_dup) lines and downregulated in the 16p11.2 deletion (16p_del) lines relative to control iPSCs (Figure 1B). Expression of pluripotency markers (Nanog and Oct4) were uniformly expressed and comparable in the 16p11.2 CNV iPSCs and control iPSCs (Supplementary Figure 1A and 1B).

Increases in gray matter volume have been associated with brain enlargement in 16p11.2 deletion carriers with macrocephaly ^24,25^. Increases in neural progenitor cell (NPC) proliferation and neuronal cell size have been associated with brain overgrowth using iPSC models of ASD with macrocephaly or 16p11.2 CNV iPSCs, respectively^26,27^. Using iPSC modeling in two- and three-dimensional systems, recent work has shown that the critical period during which gene networks become dysregulated in autism with macrocephaly occurs early at the neural stem cell stage prior to further differentiation and maturation into neurons ^28^. While sample sizes in these iPSC studies tend to be small (ranging from 3 in Deshpande et al., 2017 to 8 autistic subjects with macrocephaly in Marchetto et al., 2017 and Schafer et al., 2019), these studies nonetheless highlight that iPSCs can be used to model brain overgrowth in autism and gain insight on underlying mechanisms. To begin to model gray matter using human iPSCs, we differentiated the iPSCs into NPCs using an established NPC differentiation protocol that gives rise to cortical neurons ^29,30^. Following 12 days of directed differentiation (Figure 1C), NPCs express Pax6 mRNA at comparable and high levels of >1000-fold relative to undifferentiated iPSCs (Figure 1D). Expression of Pax6 at the protein level was also comparable across conditions (Figure 1E) and was co-expressed with nestin (Supplementary Figure 2A), an intermediate filament protein expressed in neural stem and progenitor cells^31^.

CD47, commonly known as a ‘don’t eat me’ signal, protects normal cells from getting cleared^19^, but can become overexpressed in many types of cancer cells, preventing tumorigenic cells from getting engulfed or phagocytosed ^32–34^. Recent work has shown that CD47 also plays a role in synaptic pruning^22^, suggesting that it may play an important regulatory role in neurodevelopment. We hypothesized that brain enlargement in 16p11.2 deletion carriers may be due to improper cellular elimination as a result of altered CD47 expression. To investigate this possibility, we first assessed mRNA expression levels of CD47 in the differentiated NPCs from 16p11.2 CNV carriers and control subjects. Strikingly, CD47 expression was significantly upregulated in 16p_del carriers relative to control and 16p_dup subjects (Figure 1F). Flow cytometry analysis for CD47 protein expression at the cell surface also showed a significant shift and upregulation in the mean fluorescence intensity (MFI) in 16p_del carriers relative to control and 16p_dup subjects (Figure 1G and 1H). Importantly, CD47 expression of NPCs derived from 16p_del carriers with normal size brains (average head circumference of 50.2±2.2 percentile) was comparable to control and 16p_dup individuals (Supplementary Figure 3A-E), suggesting that the upregulation of CD47 is specific to individuals with macrocephaly in 16p11.2 deletion syndrome.

With the finding of increased CD47 expression in 16p11.2 del NPCs, we hypothesized these cells would escape recognition and phagocytosis. To test the functional significance of our findings, we derived macrophages from control human blood samples and co-cultured them *in vitro* with differentiated NPCs from 16p11.2 CNV and control subjects. While NPCs derived from control and 16p_dup subjects had similar rates of phagocytosis as assessed by co-labeling with the marker CD45, there was a significant reduction in engulfment of the 16p_del NPCs (Figure 1I and 1J). Treatment with CD47 blocking antibody can effectively restore phagocytosis of cancer cells that overexpress CD47 without affecting normal cells^35–39^. To investigate whether blocking CD47 could restore phagocytic activity, we pretreated 16p_del NPCs with a CD47Ab (anti-CD47 antibody clone B6.H12) and assessed the rate of phagocytosis when co-cultured with macrophages. Strikingly, blockade of CD47 restored phagocytosis to control levels in the 16p_del NPCs (Figure 1I and 1J), potentially indicating a novel therapeutic intervention for this psychiatric disorder. Notably, CD47 expression at the mRNA and protein levels as well as phagocytosis rates were similar across groups prior to differentiation at the pluripotent stage (Supplementary Figure 1C-G), emphasizing the role of CD47 in maintaining phagocytic balance during development and highlighting that these findings were specific to the differentiated NPCs.

### Upregulation of CD47 in 16p11.2 deletion OPCs suppresses phagocytosis

In addition to increases in gray matter, brain imaging studies have shown increases in white matter volume in 16p11.2 deletion carriers with macrocephaly ^11,24,25,40,41^. While human iPSC-derived oligodendrocytes have been used to model psychiatric disorders, such as schizophrenia^24,42^, no cellular models investigating white matter overgrowth in 16p11.2 deletion syndrome have yet been developed and investigated. To begin to model white matter using human iPSCs, we differentiated the iPSCs into oligodendrocyte progenitor cells (OPCs) using an established differentiation protocol that enables iPSC-derived OPCs to engraft and function *in vivo* in the mouse brain^43^ (Figure 2A). Following 12 days of directed differentiation, the cells express pre-OPC markers, Olig2 and Nkx2.2 at the protein level^43^ (Figure 2B). With further differentiation to day 50, mRNA expression of several genes associated with the oligodendrocyte lineage (e.g. Olig1)^43^ are upregulated (Figure 2C). Following 50 days of differentiation, immunostaining and flow cytometry analyses show that the population of cells also express more mature OPC markers at the protein level, including SRY-box 10 (SOX10) (Figure 2D), the cell surface marker O4 (Figure 2E), and the chondroitin sulfate proteoglycan NG2 (Supplementary Figure 4), demonstrating that the majority of the cells are differentiating into the oligodendrocyte lineage^43^. In subsequent analyses, we collectively call the day 50 population as OPCs.

**Figure 2.**
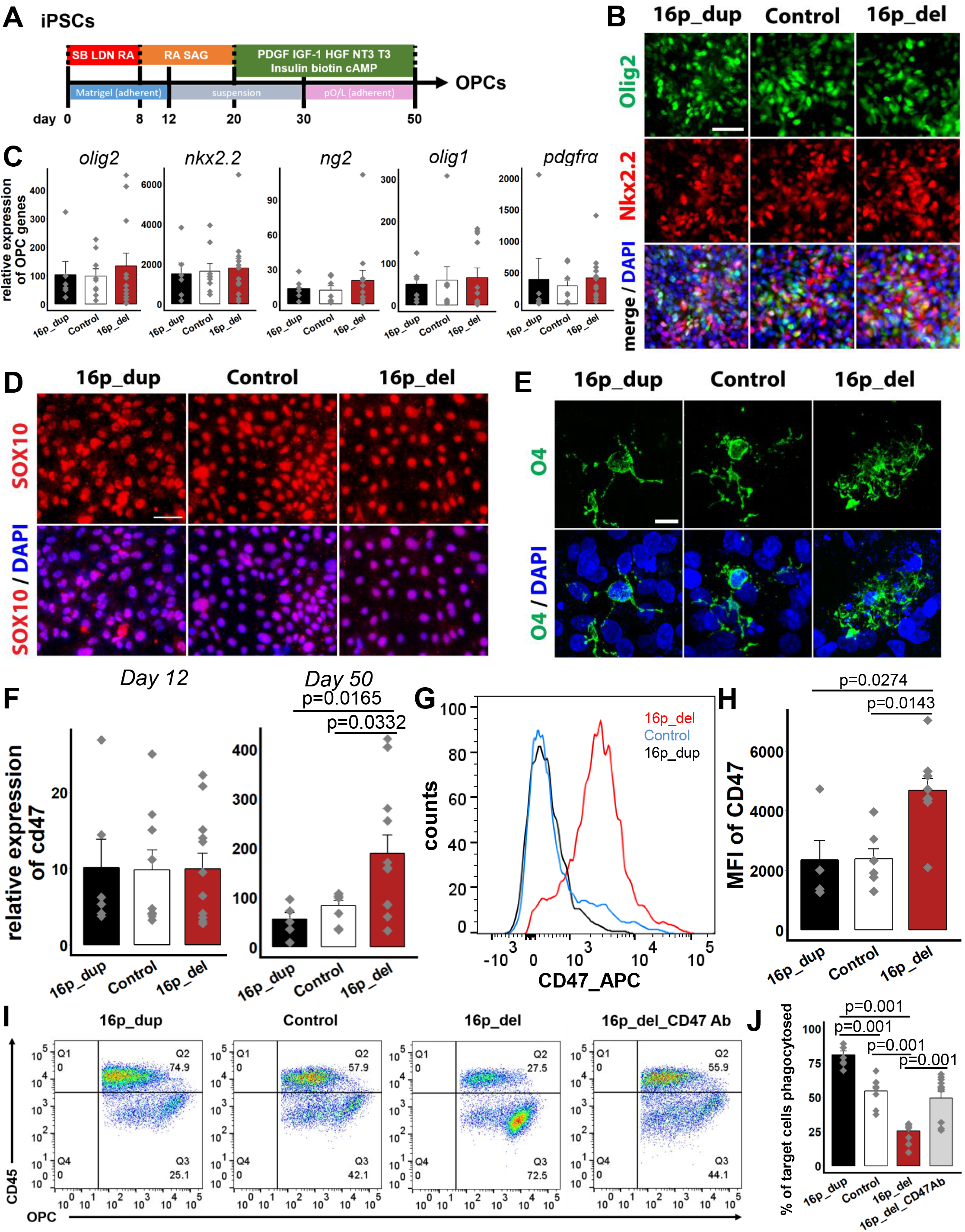
CD47 is overexpressed in 16p11.2 deletion OPCs. **A.** Schematic showing directed differentiation of iPSCs into oligodendrocyte progenitor cells (OPCs). **B.** After 12 days of directed differentiation, immunostaining shows that the cells derived from control and 16p11.2 CNV hiPSCs express pre-OPC markers Olig2 and Nkx2.2. Nuclei (DAPI, blue). **C.** Following 50 days of directed differentiation, quantitative RT-PCR shows mRNA expression levels of several genes associated with the oligodendrocyte lineage, including *olig2*, *nkx2.2*, *ng2, olig1,* and *pdgfrα* in control and 16p11.2 CNV hiPSCs. Data represent fold change relative to undifferentiated control human iPSCs. N = 3 biological replicates per cell line (16p_dup, n=2; control, n=3; 16p_del, n=4 cell lines). Mean ± SE subjected to a one-way ANOVA followed by *post-hoc* Tukey HSD Test. **D.** Immunostaining for the transcription factor, Sox10, in the 16p11.2_dup, 16p11.2_del, and control iPSC lines after 50 days of directed differentiation. **E.** Following 50 days of directed differentiation, confocal imaging shows that control and 16p11.2 CNV cells express the OPC marker O4 at the protein level. **F.** Quantitative RT-PCR results show mRNA expression levels of *cd47* in control, 16p_dup, and 16p_del lines after 12 days (left) or 50 days (right) of directed differentiation. Data represent fold change relative to undifferentiated control human iPSCs. n = 3 biological replicates per cell line (16p_dup, n=2; control, n=3; 16p_del, n=4 cell lines). **G.** Representative flow cytometry histograms showing the mean fluorescence intensities (MFI) of CD47 in OPCs after 50 days of directed differentiation. **H.** Quantification of CD47 MFI of the CD47 protein levels in OPCs derived from 16p_del lines, compared to those from control and 16p_dup lines. n = 2 biological replicates per cell line (16p_dup, n=2; control, n=3; 16p_del, n=4 cell lines). **I.** Representative flow cytometry phagocytosis plots showing rates of engulfment of OPCs when co-cultured with human derived macrophages. Phagocytosis assays were conducted using CellTrace Far Red-labeled OPCs at day 50 and human macrophages. A CD45 antibody conjugated with PE was used to specifically label human macrophages. OPCs differentiated from 16p_del lines were also pretreated with or without a CD47 blocking antibody (CD47 Ab) prior to phagocytosis assessment. **J.** The percentage of phagocytosed OPCs in 16p_del lines compared to control and 16p_dup lines. Rates of phagocytosis of OPCs in the 16p_del lines following anti-CD47 treatment with a CD47 blocking antibody (16p_del_CD47Ab) are also compared. n = 3 biological replicates per cell line (16p_dup, n=2; control, n=3; 16p_del, n=4 cell lines). All data are mean ± SE; P values were determined by one-way ANOVA followed by *post-hoc* Tukey HSD Test. Scale bar: 50 μm (**B**, **D**, **E**).

To investigate the role of CD47 in OPCs, we next assessed the mRNA expression level of CD47 following 12 days of directed differentiation, when the cells are still in the early stages of differentiation at a pre-OPC stage^43^. No significant difference in CD47 expression was detectable at this early stage across all 16p CNV and control lines (Figure 2F, left panel). However, after 50 days of differentiation, there was a significant increase in CD47 expression in the 16p_del OPCs compared with 16p_dup and control OPCs (Figure 2F, right panel). O4+ OPCs can be isolated through fluorescence activated cell sorting (FACs) using established protocols^44^. In O4+ sorted OPCs, mRNA expression of CD47 was also significantly upregulated, up to 15000-fold in 16_del relative to control and 16p_dup conditions (Supplementary Figure 5A). Across all conditions, the O4+ OPCs had high mRNA expression levels of genes associated with the oligodendrocyte lineage (Supplementary Figure 5B and 5C), low expression levels of neuronal (Tuj1) and astrocytic (GFAP) genes (Supplementary Figure 5C), and were of cortical origin as genes associated with the spinal cord had low expression (Supplementary Figure 5D). Similar to the NPCs, there was a significant shift and upregulation of the MFI of CD47 at the protein level at the cell surface of 16p_del OPCs compared to control and 16p_dup subjects after 50 days of differentiation (Figure 2G and 2H). This upregulation in CD47 corresponded with suppressed phagocytosis when the OPC populations were co-cultured with macrophages derived from control human blood samples (Figure 2I). As in NPCs, treatment with the blocking antibody, CD47Ab (anti-CD47 antibody clone B6.H12), restored phagocytosis rates to control levels in 16p_del OPCs (Figure 2I and 2J). Similar to the NPCs, CD47 expression of OPCs derived from 16p_del carriers with normal size brains (average head circumference of 50.2±2.2 percentile) was comparable to control and 16p_dup individuals (Supplementary Figure 6A-C), providing further support that the upregulation of CD47 is specific to individuals with macrocephaly in 16p11.2 deletion syndrome.

### Overexpression of CD47 leads to suppressed phagocytosis of 16p11.2 deletion NPCs and OPCs *in vivo*

Thus far, our results show that overexpression of CD47 in the 16p_del NPCs and OPCs is associated with suppressed phagocytosis *in vitro*. We next investigated whether phagocytosis rates would also be altered and recapitulated in an *in vivo* setting following intracerebral injections of 16p11.2 CNV-derived NPCs or OPCs. To do so, following *in vitro* directed differentiation, hiPSC-derived NPCs and OPCs were labeled with CellTrace red fluorescent dye and injected intracerebrally into individual NOD-scid IL2gammanull (NSG) P2 pups (Figure 3A). 24h after injection, pups were euthanized and brains were dissociated for flow cytometry analyses of phagocytosis. Within the target cell population identified by CellTrace Far Red, the percentage of cells co-labeled with CD45 and CD11b, markers of activated microglia and infiltrating macrophages, were analyzed to determine phagocytosis rates (Figure 3A)^45^. Similar to the *in vitro* experiments, the percentages of engulfed or phagocytosed NPCs (Figure 3B and 3C) and OPCs (Figure 3D and 3E) were significantly suppressed *in vivo* in the 16p_del conditions relative to control and 16p_dup conditions. Strikingly, treatment with an anti-CD47 blocking antibody of 16p_del NPCs and OPCs prior to intracerebral injections restored *in vivo* phagocytosis rates to control levels (Figure 3B-E).

**Figure 3.**
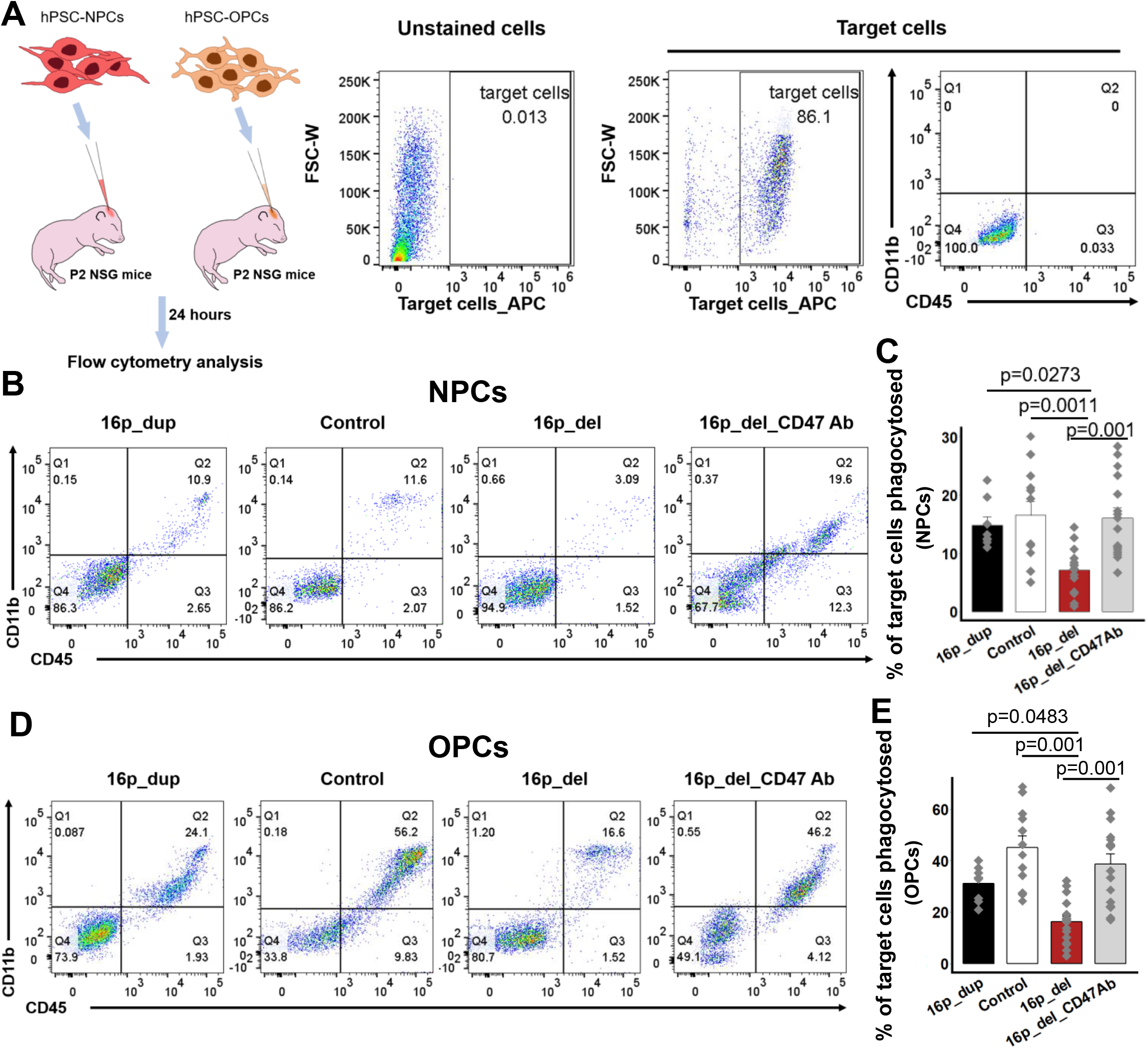
Overexpression of CD47 suppresses *in vivo* phagocytosis of 16p 11.2 deletion NPCs and OPCs. **A.** Schematic of *in vivo* phagocytosis assay using NPCs and OPCs differentiated from control, 16p11.2 deletion (16p_del), and 16p11.2 duplication (16p_dup) iPSC lines. NPCs and OPCs differentiated from 16p_del lines were also pretreated with or without a CD47 blocking antibody prior to phagocytosis assessment. CellTrace Far Red-labeled 12 day differentiated NPCs or 50 day differentiated OPCs were injected into P2 NSG mice brain. 24 hours post injection, the brains were isolated and disassociated for flow cytometry analysis. Unstained cells show negative expression of CellTrace Far Red-labeled target cells (NPCs or OPCs). Mouse-CD11b-FITC and mouse-CD45-PE were used to detect mouse microglia and infiltrating macrophages within the population of CellTrace Far Red-labeled NPC- or OPC-target cells. Within the population of target cells, CD11b and CD45 expression remains negligible prior to *in vivo* injection. **B.** Representative flow cytometry phagocytosis plots showing rates of engulfment of NPCs 24 hours post injection into NSG pup brains. Phagocytosed NPCs are indicated by the co-expression of CD11b and CD45 (Q2). **C.** The percentages of phagocytosed NPCs in 16p_del relative to control and 16p_dup conditions. Rates of phagocytosis when 16p_del NPCs are pre-treated with a CD47 blocking antibody (16p_del_CD47Ab) are also shown. **D.** Representative flow cytometry phagocytosis plots showing rates of engulfment of OPCs 24 hours post injection into NSG pup brains. Phagocytosed OPCs are indicated by the co-expression of CD11b and CD45 (Q2). **E.** The percentages of phagocytosed OPCs in 16p_del lines relative to control and 16p_dup lines. Rates of phagocytosis when 16p_del OPCs are pre-treated with a CD47 blocking antibody (16p_del_CD47Ab) are also shown. All data are mean ± SE. n=2 pups per cell line, 2 technical replicates per pup (16p_dup, n=2; control, n=3; 16p_del, n=4 cell lines); P values were determined by one-way ANOVA followed by *post-hoc* Tukey HSD Test.

### Anti-CD47 treatment restores *in vivo* phagocytosis of 16p11.2 deletion NPCs and OPCs

To assess therapeutic implications, we next evaluated whether intraperitoneal (*i.p.*) anti-CD47 treatment could restore phagocytosis of 16p11.2_del NPCs and OPCs *in vivo*, analogous to an approach that has proven to be beneficial in human clinical trials of patients with lymphoma^39^. After intracerebrally injecting CellTrace Far Red-labeled 16p11.2_del NPCs and OPCs into NSG pups, we treated the pups with anti-CD47 (Hu5F9-G4) or human IgG control antibodies intraperitoneally and analyzed phagocytosis by flow cytometry after 24h (Figure 4A). Compared to IgG control, treatment with anti-CD47 antibody was associated with a marked increase in phagocytosis, both in 16p_del NPCs (Figure 4B and 4C) and 16p_del OPCs (Figure 4D and 4E). Additionally, we used antibodies to ionizing calcium-binding adaptor molecule (Iba1), a microglial and macrophage-specific calcium-binding protein that is involved in membrane ruffling and phagocytosis^46^, to immunolabel and identify phagocytic cells in NSG pup brain tissue sections. Consistent with the flow cytometry analyses, increased Iba1+ cells engulfing target cells marked by the human cytoplasmic marker (STEM121) along with CellTrace Far Red at the injection site were evident in 16p_del brains treated with anti-CD47 antibody relative to control IgG treatment (Figure 4F). Taken together, these results indicate that CD47-blockade could act as a therapeutic agent in clearing NPCs and OPCs in the 16p11.2 deletion syndrome.

**Figure 4.**
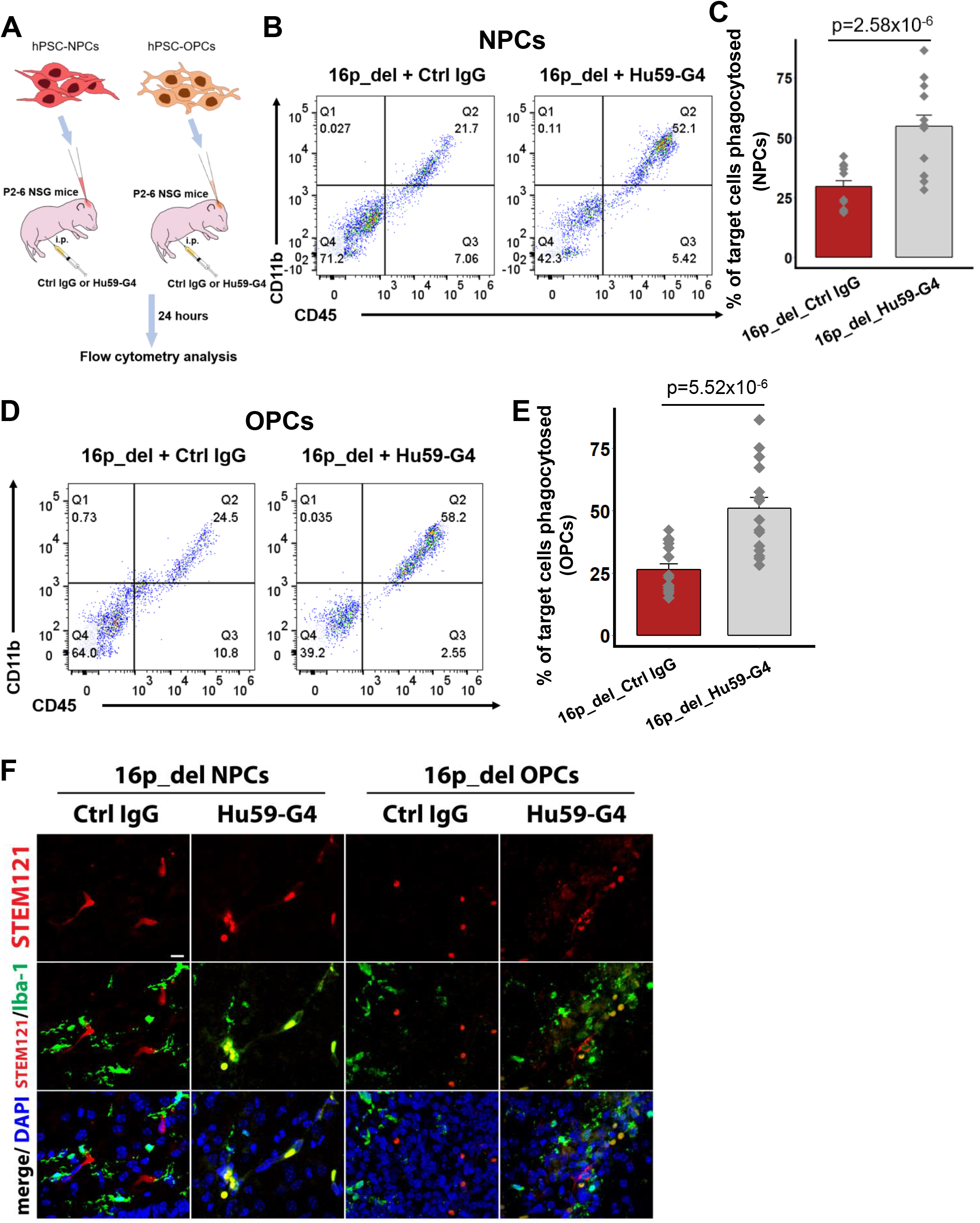
Anti-CD47 treatment restores phagocytosis of 16p11.2 deletion NPCs and OPCs *in vivo*. **A.** Schematic of *in vivo* phagocytosis assay using NPCs and OPCs differentiated from 16p11.2 deletion (16p_del) iPSC lines and treated with human IgG control or anti-CD47 antibody (Hu59-G4) intraperitoneally (IP). CellTrace Far Red-labeled 12 day differentiated NPCs or 50 day differentiated OPCs were injected into P2-P6 NSG mice brain with IP treatment of human IgG control or anti-CD47 antibody. 24 hours post injection, the brains were isolated and disassociated for flow cytometry analysis. Mouse-CD11b-FITC and mouse-CD45-PE were used to detect mouse microglia and infiltrating macrophages within the population of CellTrace Far Red-labeled NPCy- or OPC-target cells. **B.** Representative flow cytometry phagocytosis plots showing rates of engulfment of NPCs. Phagocytosed NPCs are indicated by the co-expression of CD11b and CD45 (Q2). **C.** Percentages of phagocytosed NPCs derived from 16p_del iPSCs in control IgG-treated or Hu59-G4-treated pups. **D.** Representative flow cytometry phagocytosis plots showing rates of engulfment of OPCs. Phagocytosed OPCs are indicated by the co-expression of CD11b and CD45 (Q2). **E.** Percentages of phagocytosed OPCs derived from 16p_del iPSCs in control IgG- treated or Hu59-G4-treated pups. **F.** Confocal images of CellTrace Far Red-labeled NPCs or OPCs in NSG pup brains probed for the human cytoplasmic marker STEM121 (red) and Iba1 (green) for phagocytic cells, and nuclei (DAPI, blue) in control IgG or Hu59-G4-treated mice. Scale bar, 50 μm. All data are mean ± SE. n=2 pups per cell line, 2 technical replicates per pup (16p_dup, n=2; control, n=3; 16p_del, n=4 cell lines); P values were determined by two-tailed student’s *t*-test.

### Upregulation of the pro-phagocytic signal Calreticulin in 16p11.2 deletion NPCs and OPCs

Under normal conditions, cell homeostasis is maintained by the balance of pro- and anti-phagocytic signals. Calreticulin is a dominant pro-phagocytic signal (commonly known as an ‘eat me’ signal) that is upregulated on the surfaces of cancer cells as well as damaged and aged cells, indicating to the phagocytic system that the cells should be eliminated^47^. However, due to high levels of CD47, which dominates the pro-phagocytic calreticulin signal, these cells fail to be eliminated^20^. Importantly, translocation of calreticulin to the cell surface of target cells determines cell removal^47^. Here, we assessed cell surface levels of calreticulin (CRT) by flow cytometry in the differentiated NPCs and OPCs. NPCs and OPCs derived from 16p_del iPSCs had a significant upregulation of CRT relative to control and 16p_dup subjects (Figure 5A-B, 5D-E). Furthermore, CRT binds to asialoglycans on the surface of cells, therefore the more potential binding sites, desialated glycans, the more potential calreticulin there is to promote interrogation and clearance by the phagocytic system. The absence of sialic acid on glycans can be determined by using a lectin, Phaseolus Vulgaris Leucoagglutinin (PHA-L)^47^. Staining with PHA-L on NPCs and OPCs derived from control and 16p_CNV iPSCs shows a marked increase in PHA-L binding in 16p_del subjects (Figure 5C and 5F), indicating increased presence of asialoglycan binding sites for CRT. In individuals with 16p11.2 deletion with normal head circumferences (no macrocephaly), expression of CRT was comparable to control subjects (Supplementary Figure 3F, Supplementary Figure 6D). Together, this indicates that the NPCs and OPCs in 16p11.2 deletion syndrome with brain enlargement should be eliminated but fail to be due to high expression of CD47. These data further support the importance of anti-CD47 treatment as a means to promote clearance of unhealthy NPCs and OPCs in 16p11.2 deletion syndrome with brain overgrowth.

**Figure 5.**
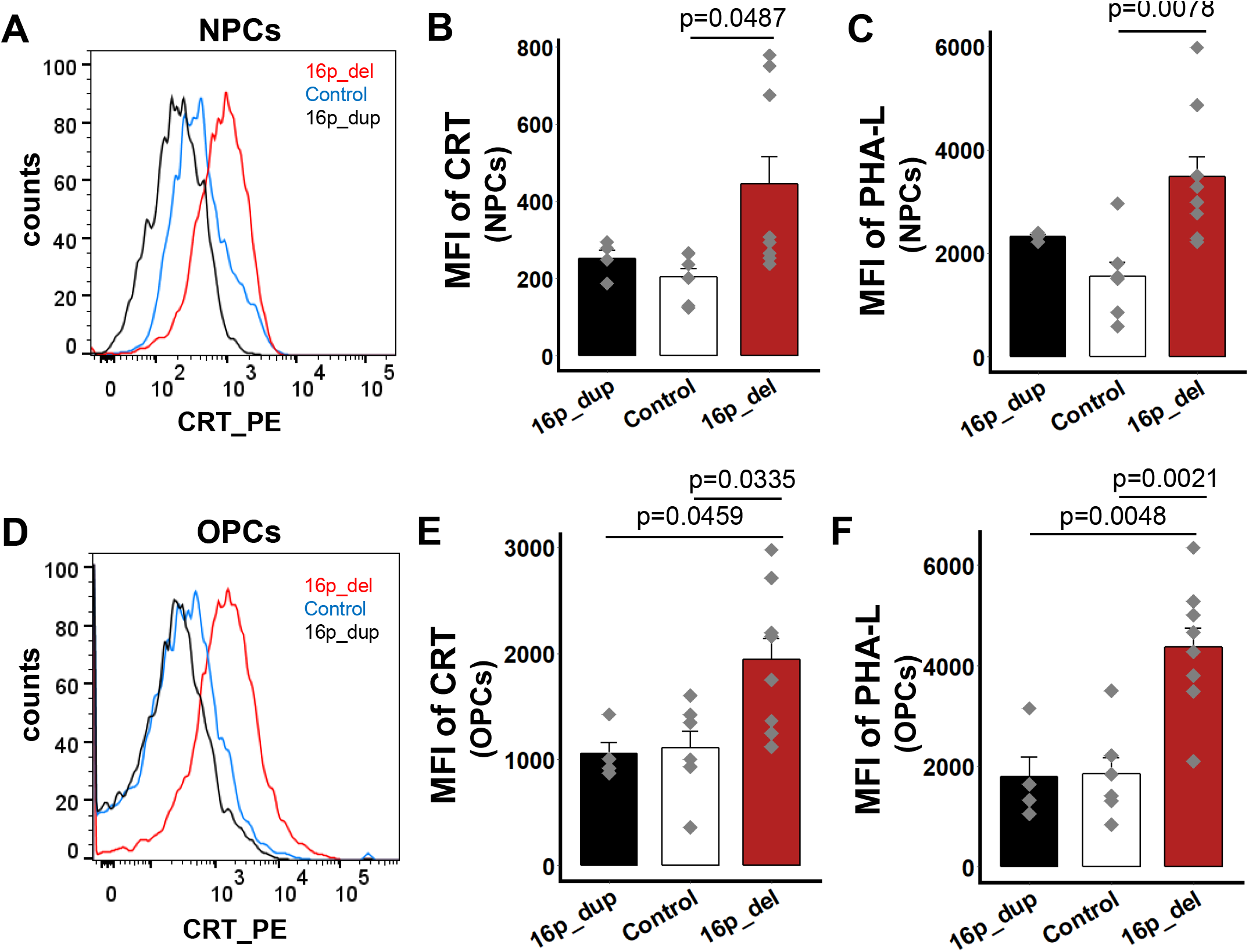
Calreticulin, a pro-phagocytic signal, is upregulated in 16p11.2 deletion NPCs and OPCs. **A.** Representative histograms show the mean fluorescence intensities (MFI) of cell-surface expression of Calreticulin (CRT) in NPCs derived from control, 16p_del, and 16p_dup lines. **B.** Quantification of CRT MFI of cell-surface expression of CRT in the 16p_del NPCs relative to 16p_dup and control NPCs. **C.** Quantification of Phaseolus Vulgaris Leucoagglutinin (PHA-L), indicative of asialoglycan binding sites for CRT, in NPCs in 16p_deletion lines relative to control and 16p_duplication lines. **D.** Representative histograms show the MFI of cell-surface expression of CRT in OPCs derived from control, 16p_del, and 16p_dup lines. **E.** Quantification of CRT MFI of cell-surface expression of CRT in the 16p_del OPCs relative to 16p_dup and control OPCs. **F.** Quantification of PHA-L in OPCs in 16p_deletion lines relative to control and 16p_duplication lines. n=2 biological replicates per cell line (16p_dup, n=2; control, n=3; 16p_del, n=4 cell lines). All data are mean ± SE; P values were determined by one-way ANOVA followed by *post-hoc* Tukey HSD Test.

## Discussion

While CD47 has been extensively studied in cancer^20,32–35,38^, this study begins to uncover its implications for neurodevelopmental disorders associated with brain overgrowth. Recent work has shown that CD47 plays a critical role in regulating synaptic pruning during early brain development^22^, providing further support that the same pathways can go awry in neurodevelopmental disorders. While a number of studies are beginning to show changes in NPC proliferation^26,28^ and neuronal size^27^ associated with autism with brain enlargement, these studies focused on changes in gray matter and have only investigated molecular and cellular mechanisms in cultures *in vitro*^26–28^. We demonstrate that CD47 is upregulated in two distinct cell types (NPCs and OPCs) at particular stages of development in 16p11.2 deletion syndrome, leading to suppressed phagocytosis both *in vitro* and *in vivo*. Moreover, we provide evidence that these findings are specific to individuals with macrocephaly and not those with normal head circumferences with 16p11.2 deletion syndrome. We demonstrate that anti-CD47 treatment can act as a therapeutic agent for clearing unhealthy cells in cellular and mouse models, potentially indicating that these forms of therapy could be translated to selected autistic individuals with brain overgrowth early in the disease.

In acute myeloid leukemia (AML), the hematopoietic stem cell precursors of AML out-compete normal hematopoietic stem cells, enabling the leukemia stem cells to expand beyond by countering signaling associated with programmed cell removal, such as calreticulin, with overexpression of CD47^20,48–50^. The imbalance between CD47 and calreticulin observed in this study may indicate that specific NPCs and OPCs in the 16p deletion syndrome with brain overgrowth undergo clonal expansion and outcompete normal NPCs and OPCs. Future work should evaluate whether expansion of CD47hi NPC or OPC clones contribute to brain overgrowth in psychiatric and neurodevelopmental disorders, such as autism. Interestingly, a surprising number of genes (e.g. PTEN, CHD8) and cellular pathways involved in cancer have been found to overlap with those implicated in autism^51–54^. This not only indicates potentially shared mechanisms between autism and cancer, but also that the breadth of therapeutic agents identified for cancer could be beneficial in autism and other neurodevelopmental disorders. Here, we demonstrate the efficacy of anti-CD47 monoclonal antibodies as a potential therapeutic in restoring phagocytosis of 16p11.2 deletion NPCs and OPCs. Within the 16p11.2 locus, approximately 29 genes are affected in 16p11.2 CNV carriers^5^. In future work, it would be interesting to investigate which of these 29 genes directly regulates the CD47 pathway and/or the regulation of asialoglycoproteins (binding sites for calreticulin), which could have important implications for cancer, neurodevelopment, and other conditions associated with CD47 signaling. Similarly, as in some forms of cancer^55^, it would be interesting to investigate whether changes in epigenetics (e.g. H3K27 acetylation) of 16p11.2 deletion NPCs and OPCs influence enhancers and super enhancers to regulate CD47 expression.

We acknowledge that while the sample sizes in this study are small (but comparable to other iPSC studies modeling subtypes of psychiatric disorders)^26–28,56^, they would benefit from including more patients with 16p11.2 deletion and/or other disorders with defined macrocephaly. Validating these findings in postmortem brain tissue in 16p11.2 deletion individuals with brain overgrowth would also be valuable, though this resource is limited and constrained by the fact that most brain banks worldwide do not have head circumference or brain volume data tagged with postmortem human brain tissue. Nonetheless, iPSCs are increasingly proving to be a valuable resource in understanding the mechanisms regulating brain development and growth. In recent work, proliferation of NPCs has been shown to be suppressed in human iPSC models of patients with psychiatric disorders and reduced brain volume with 16p13.11 microduplication^56^. Investigating whether imbalances between pro-and anti-phagocytic signals, such as calreticulin and CD47, regulate brain size more generally in other conditions associated with brain overgrowth or undergrowth may lead to novel therapeutic strategies to prevent and treat these neurodevelopmental disorders.

## Materials and Methods

### Characterization and maintenance of hiPSCs

Induced pluripotent stem cell (iPSC) lines used in this study are listed in Supplementary Table 1 and obtained from the Simons Foundation Autism Research Initiative (SFARI) database *via* Simons Variation in Individuals Project (VIP). Full phenotypic information and clinical information for each patient were provided by VIP. A total of eleven human iPSC lines were used in the study. Cells were cultured and maintained in mTeSR medium (Stem Cell) with 10 μM ROCK inhibitor (Y27632, Stemgent) at 37°C and 5% CO_2_ on Matrigel (BD Biosciences). Prior to differentiation, all hiPSC lines were immunostained against OCT3/4 and NANOG for pluripotency analysis. We also verified that genes within the 16p11.2 locus of the iPSCs have altered gene expression corresponding to either the 16p11.2 deletion or duplication. Control lines have been confirmed to have no genetic abnormalities.

### Regulatory and Institutional Review

All human pluripotent stem cell experiments were conducted in accord with experimental protocols approved by the Stanford Stem Cell Research Oversight (SCRO) committee.

### Generation of neural progenitor cells (NPCs)

Control and 16p11.2 CNV iPSCs were differentiated into cortical neural progenitor cells (NPCs) as previously described^29,30^. The iPSC cultures were plated at a high-density monolayer onto GelTrex (ThermoFisher) coated wells. When cells were confluent, neuroectoderm differentiation began in Essential 6 Media (ThermoFisher) supplemented with small chemical inhibitors of the TGFβ, SMAD, and Wnt pathways, 500 nM LDN193189 (Tocris) + 10 μm SB431542 (Tocris) + 2 μM of XAV939 (Tocris) for three-days and then 500 nM LDN193189 + 10 μm SB431542 for the remaining nine days of differentiation. Media was changed daily.

### Directed differentiation of oligodendrocyte progenitor cells (OPCs)

Oligodendrocyte progenitor cells were differentiated from control and 16p11.2 CNV iPSC lines following published protocols^43^. In brief, iPSCs were plated at a density of 1 × 10^5^ per well on Matrigel-coated six-well plates containing mTeSR and 10 μM ROCK inhibitor. The differentiation medium containing DMEM/F-12 (Life Technologies), 10 μM SB431542 (Stemgent), 250 nM LDN193189 (Stemgent), 100 nM all-trans retinoic acid (RA, Sigma-Aldrich), 1 μM smoothened agonist (SAG, EMD Millipore) was applied for 12 days. Fresh media was changed daily. Subsequently, cells were detached using a Corning cell lifter (Sigma-Aldrich) and transferred to ultra-low-attachment plates for the formation of spheres. From day 12, the differentiation media was changed every other day. OPC differentiation was induced in the OPC induction media containing DMEM/F-12, 1× MEM non-essential amino acids (NEAA) solution (Life Technologies), 1× GlutaMAX (Life Technologies), 2-mercaptoethanol 1× (Life Technologies), Penicillin/Streptomycin (PenStrep; Life Technologies), 1× N-2 supplement (Life Technologies), B27 Supplement without VitA (Life Technologies), 10 ng/ml Recombinant human PDGF-AA, CF (R&D Systems), 10 ng/ml Recombinant human IGF-I, CF (R&D Systems), 5 ng/ml Recombinant human HGF (R&D Systems), 10 ng/ml Neurotrophin 3 (NT3; EMD Millipore), 25 μg/ml Insulin solution, human (Sigma-Aldrich), 100 ng/ml Biotin (Sigma-Aldrich), 1 μM N6,2′-O-Dibutyryladenosine 3′,5′-cyclic monophosphate sodium salt (cAMP; Sigma-Aldrich) and 60 ng/ml 3,3,5-Triiodo-l-thyronine (T3; Sigma-Aldrich). On day 30, aggregates were plated for attachment on poly-l-ornithine/laminin-coated plates. The oligodendrocytes were continued to be expanded and differentiated in the OPC induction medium until harvested. The cells were immunostained with oligodendroglial specific markers, such as Olig2, Nkx2.2, and O4, to confirm their progression into the oligodendrocyte lineage.

### Immunocytochemistry

Cells were fixed in cold 4% paraformaldehyde in phosphate buffered saline (PBS; Santa Cruz Biotechnology) for 20 minutes to one hour under room temperature. Subsequently, the cells were rinsed three times in PBS. For the nuclear staining, cells were permeabilized with 0.3% Triton X-100 (Fisher Bioreagents) for 15 minutes and washed three times with PBS. Subsequently, blocking solution containing 5% normal donkey serum (Jackson ImmunoResearch) /0.3 % TrionX-100 in PBS was applied for one hour. The primary antibodies and corresponding dilutions used were: rabbit-anti-PAX6 (1:500; Biolegend); mouse-anti-Nestin (1:500; R&D Systems); rabbit-anti-Olig2 (1:100; EMD Millipore); mouse-anti-Nkx2.2 (1:50; DSHB); goat-anti-Sox10 (1:500; R&D Systems); mouse-anti-Oct3/4 (1:500; SantaCruz Biotechnology) and rabbit-anti-NANOG (1:500; Stemgent). After overnight primary antibody incubation at 4°C, cells were rinsed in PBS/0.3 % TrionX-100, incubated with fluorescently tagged secondary antibodies (Alexa-Fluor goat/donkey-anti-primary antibody species IgG 488 or 549; Life Technologies) for one hour. Nuclear dye, DAPI (4,6-diamidino-2-phenylindol, Life Technologies), was used to stain all cells. All images were acquired using a Leica Fluorescent Microscope.

The mouse-anti-O4 surface antigen (1:500; clone 81; EMD Millipore) was used to identify hiPSC-derived OPCs. After 15 minutes of fixation in cold 4% paraformaldehyde/PBS, the cells were blocked in 1% BSA (Sigma-Aldrich) /5% normal donkey/ PBS for one hour at room temperature, and subsequently immunolabeled as described above. The images were taken by confocal microscopy.

### RNA isolation and quantitative real-time PCR

Total RNA was isolated from control and 16p11.2 CNV hiPSCs, and hiPSC-derived NPCs and OPCs using the RNeasy Mini Kit (QIAGEN) following manufacturer’s instructions. RNA was quantified using the NanoDrop spectrophotometer. Reverse transcription was performed subsequently to synthesize cDNA using SuperScript IV VILO Master Mix with ezDNase (Thermo Fisher) using the same quantity of RNA for all samples.

Quantitative Real-Time PCR (qRT-PCR) was performed using the SYBR green system. One reaction included SYBR green mix (Applied Biosystems), the forward and reverse target gene primers, and 100–120 ng cDNA. The StepOnePlus Real-Time PCR System (AppliedBiosystems) was used to run the qRT-PCR experiment. The expression of RNA transcripts was analyzed using the ΔΔCT method. Three biological, as well as three technical replicates, were conducted. All samples were normalized to the expression of the housekeeping gene *gapdh*. Primer sequences used in this study are listed in Supplementary Table 2.

### Confocal microscopy

O4 staining in OPCs derived from control and 16p11.2 CNV iPSC lines, and the mouse brain tissue sections stained with the monoclonal Human Cytoplasmic Marker (STEM121) and microglial/macrophage marker (Iba1) were imaged using Zeiss LSM710 Confocal Microscope with excitation laser lines at 405 (DAPI), 488 (GFP) and 594 nm using a 40X plan-fluor oil-immersion objective.

### Flow Cytometry Analysis

The iPSCs, NPCs, or OPCs were dissociated using Accutase (Life Technologies). The target cells were subsequently suspended in FACS buffer containing 2% bovine serum albumin (BSA; Sigma), and 2 mM EDTA in PBS. Cells were then stained with following antibodies for 30 minutes on ice: Anti-human-NG2 Chondroitin Sulfate Proteoglycan, Cy3 (1:100; Millipore); Anti-human-CD47-APC (Clone: B6H12; 1:100; ThermoFisher); anti-human-Calreticulin-PE (1:100; Enzo Life Sciences); anti-mouse/human-CD11b-FITC (Clone: M1/70; 1:100; Biolegend); anti-human-CD45-PE (Clone: 2D1; 1:100; Biolegend); and anti-mouse-CD45-PE (Clone: 30-F11; 1:100; Biolegend). The lectin fluorescein labeled Phaseolus Vulgaris Leucoagglutinin (PHA-L) (1:200; Vector Laboratories) was used to stain for PHA-L. For O4 staining, the cells were first incubated in primary antibody mouse-anti-O4 antibody (Clone 81; 1:100; EMD millipore), followed by secondary antibody incubation, anti-Mouse IgM Alexa Fluor 488 (1:1000; Life Technologies). The cells were then washed twice with FACS buffer.

DAPI (Sigma) was used to exclude dead cells. Immediately prior to flow cytometric analysis, the cells were strained through a 100 μM filter. Flow cytometry analyses were performed on the BD FACS Aria II (Becton Dickinson) and data were analyzed using FlowJo software (FloJo LLC). CD47 expression was measured using mean fluorescence intensity (MFI). The MFI of cell surface calreticulin (CRT) expression was obtained from the CD47+ population. Representative unstained controls for CD47 and cell surface CRT MFI are shown in Supplementary Figure 7. The MFI for PHA-L expression was also measured.

### O4-positive cell sorting

To obtain a pure population of O4-expressing OPCs, the mouse-anti-human-O4 antibody (Clone 81; 1:100; EMD millipore) was used to stain the 50 day-hiPSC derived-OPCs. The cells were incubated in the primary antibody for 40 minutes on ice, washed with FACS buffer, and subsequently stained with Alexa Fluor 488 goat anti-mouse IgM (μ-chain; Life Technologies) for 30 minutes on ice. The cells were rinsed twice with FACS buffer. DAPI stain was used to exclude dead cells. Immediately prior to flow cytometric analysis, the cells were strained through a 100 μM filter.

O4-positive cell sorting were performed on the BD FACS Aria II (Becton Dickinson) using Accudrop experiments. Cells were sorted into a clean FACS tube containing Buffer RLT (RNeasy Mini Kit, QIAGEN) for RNA extractions as described above. For each cell line, 1 × 10^4^ cells were collected.

### *In vitro* phagocytosis and anti-CD47 treatment

To generate human peripheral blood-derived macrophages, CD14+ monocytes were enriched from human peripheral blood (purchased from Stanford Blood Center) and cultured in Iscove’s Modified Dulbecco’s Medium (IMDM; ThermoFisher) with 10% human serum to differentiate to macrophages. After 7 days differentiation, human macrophages are ready for the phagocytosis. The hiPSCs and hiPSC-derived NPCs (at 12 days differentiation) and OPCs (at 50 days differentiation) were fluorescently labeled with CellTrace (far red; ThermoFisher). The human macrophages were lifted using TrypLE (Gibco) and Corning cell lifter, and transferred to 10% human serum/DMEM. In a 3:1 ratio of macrophages (approx. 7.5 × 10^4^ cells) to target cells (approx. 2.5 × 10^4^ cells), the cells were mixed in sterile polystyrene tubes (Fisher Scientific), and incubated at 37 °C for 2 hours on a shaker (250 rpm) to promote macrophage and target cell interaction. Subsequently, the cells were washed in cold FACS buffer (2 mM EDTA/2% BSA/PBS) and stained for human marker Human-CD45-PE (Clone: 2D1; 1:100; Biolegend) before analysis on the FACS Aria II machine. 7-AAD (Biolegend) stain was used to exclude dead cells.

Treatment with the CD47 blocking antibody was performed immediately after CellTrace labeling. The *InVivo*MAb anti-human CD47 antibody (Clone: B6.H12; BioXcell) was diluted to 10 μg/ml in DMEM and applied to the target cells for 30 minutes on ice. Samples were then washed in 10%FBS/DMEM and processed for co-incubation with human macrophages as described above.

### Mice Maintenance

The NOD scid gamma mice (NSG mice) were kept at a high barrier conditions and under pathogen-free environment in Lokey Stem Cell Building at Stanford University. All animal handling, surveillance, and experimentation were performed in accordance with approval from the Stanford University Administrative Panel on Laboratory Animal Care.

### *In vivo* phagocytosis and anti-CD47 treatment

The target cells were detached using Accutase, counted, stained following manufacturer’s recommendation with CellTrace (far red; ThermoFisher), and stored on ice until injection. Each P2-6 NOD-scid IL2gammanull (NSG) mouse was anesthetized by hypothermia and received a bilateral injection of 1μL (0.5μL DMEM/F12 and 0.5μL HBSS) containing 30,000 cells of either NPCs or OPCs using a 10μL Hamilton microsyringe (Hamilton Company). The injection site was determined by injecting at midline, +1mm rostral of lambda and injection took place through the skin and soft skull. The pups were gently warmed and then returned back to their mother when active. Twenty- four hours post-injection, pups were euthanized by decapitation under hypothermia and the fresh brains removed and placed in cold HBSS media on ice as previously described^57^. 2mm × 2mm cortical piece of tissue from the midline injection site was subject to neural dissociation (Neural Tissue Dissociation Kit; Miltenyi Biotec). Following dissociation, the cells were strained through a 100μm filter and suspended in FACS buffer. All cells were then stained using Mouse/Human-CD11b-FITC (Clone: M1/70; 1:100; Biolegend); Mouse-CD45-PE (Clone: 2D1; 1:100; Biolegend). Dead cells were excluded using DAPI, and samples were analyzed on FACS Aria II machine. The targeted NPCs and OPCs population could be determined by APC-positive gating. The relative ratio of phagocytosed target cells was determined by co-expression of CD11b-FITC- and CD45-PE-positive cells.

Treatment with the CD47 blocking antibody was performed immediately after cell injection. 1μl of 250μg/mL of CD47 blocking antibody (Hu5F9-G4, acquired from Forty Seven) was injected intraperitoneally into the pups. The pups were gently warmed and then returned back to their mother when active. Twenty-four hours post-injections, brains were processed, stained, and analyzed by flow cytometry as described above.

### Serial Sectioning and Staining

For histological evaluations, brains were extracted and placed in 4% Paraformaldehyde (PFA; Electron Microscopy Science) for 24 hours at 4 °C. Brains were then placed in 30% sucrose solution (Sigma) dissolved in deionized water for 48 hours and/or until brains sank to the bottom. They were then embedded in optimal cutting temperature compound (OCT; Fisher Scientific). Serial coronal sections were cut at 20 μm thick using a Cryostat at 200μm apart for a total of 5 sections per animal spanning the injection site and then stored at −80 °C. At room temperature (RT), the dried sections were postfixed with 10% formalin, washed 3 times in 1x PBS, and blocked in 2% donkey serum with 0.1% Triton X-100 for 1 h. Slides were then incubated with primary antibody diluted in blocking solution at 4 °C overnight: mouse-monoclonal Human STEM121 (1:500; Takara) for detecting human cytoplasmic protein and Rabbit-anti-Iba1 (1:500; Wako) for identifying mouse microglia and infiltrating macrophages. Sections were washed 5 times with 1 × PBS and incubated with fluorescently tagged secondary antibodies (Alexa-Fluor mouse IgG 488 or rabbit 594; Life Technologies) for one hour at RT. Following staining, the slides were then washed 3 × in 1 × PBS and then mounted in Pro-long anti-fade mounting solutions with DAPI.

### Statistical analysis

All data are graphed as bar blots using the ggplot2 library in RStudio Desktop 1.1.463 environment (https://www.statmethods.net/advgraphs/ggplot2.html). Mean values with standard errors of the mean (SEM) are reported. Student’s unpaired two-tailed *t test* was used for comparisons of two experimental groups. Multiple comparisons were analyzed using one-way analysis of variance (ANOVA) followed by *post-hoc* Tukey HSD Test. P value ≤ 0.05 was considered statistically significant. When p values are not reported in the figures, there was no statistical significance.

## Acknowledgements

We are grateful to all of the families at the participating Simons Variation in Individuals Project (Simons VIP) sites, as well as the Simons VIP Consortium. We appreciate obtaining access to phenotypic data on SFARI Base. We thank the Flow Cytometry Core facilities for their assistance. We are very grateful to Rahul Sinha, Danielle Sambo, Kyle Loh, Liang Ma, Mellisa Stafford, Nobuko Uchida, and Theo Palmer for their assistance and feedback on the manuscript and David Amaral, Joachim Hallmayer, Ruth O’Hara, Christine Nordahl, and members of the Weissman laboratory for helpful discussions. Diversity supplement R01CA086065 (to KM). This work was supported by grants from the Stanford University School of Medicine and a Siebel Fellowship awarded to S.C..

## Data availability

All data generated or analyzed are during this study are included in this published article (and its supplementary information files).

---

**Supplementary Figure 1. CD47 expression is not altered at the pluripotency stage in 16p11.2 CNV iPSC lines.**

**A.** Quantitative RT-PCR for mRNA expression levels of pluripotency genes in the 16p_dup, 16p_del, and control iPSC lines. Data represent fold change relative to undifferentiated control human embryonic stem cells (hESCs). n = 3 biological replicates per cell line (16p_dup, n=2; control, n=3; 16p_del, n=4 cell lines). **B.** Immunostaining for the pluripotency markers, Nanog and Oct4, in the 16p11.2 CNV and control iPSC lines. Scale bar: 50 μm. **C.** Quantitative RT-PCR of *cd47* mRNA expression at the iPSC stage in 16p_dup, 16p_del, and control iPSCs. Data represent fold change relative to undifferentiated control human iPSCs. n = 3 biological replicates per cell line (16p_dup, n=2; control, n=3; 16p_del, n=4 cell lines). **D.** Representative histograms showing the mean fluorescence intensities (MFI) of CD47 at the iPSC stage in 16p11.2 CNV and control iPSCs. **E.** Quantification of CD47 MFI in 16p11.2 CNV and control iPSCs. n = 2 biological replicates per cell line (16p_dup, n=2; control, n=3; 16p_del, n=4 cell lines). **F.** Representative flow cytometry phagocytosis plots showing rates of engulfment of iPSCs when co-cultured with human derived macrophages. Phagocytosis assays were conducted using CellTrace Far Red-labeled undifferentiated iPSCs and human macrophages. A CD45 antibody conjugated with PE was used to specifically label human macrophages. **G.** The percentages of phagocytosed iPSCs compared across 16p_dup, 16p_del, and control iPSC lines. n = 3 biological replicates per cell line (16p_dup, n=2; control, n=3; 16p_del, n=4 cell lines). All data are mean ± SE and were subjected to a one-way ANOVA followed by *post-hoc* Tukey HSD Test.

**Supplementary Figure 2. Directed differentiation of 16p11.2 CNV and control iPSC lines into neural progenitor cells**.

**A.** Immunostaining for the cortical neural progenitor marker, Pax6 (green) and pan- neural marker Nestin (red), in the 16p11.2_dup, 16p11.2_del, and control lines after 12 days of differentiation into NPCs. DAPI (blue) used to stain nuclei. Scale bar: 50 μm.

**Supplementary Figure 3. CD47 expression is not altered in NPCs derived from 16p11.2 deletion lines without macrocephaly**.

**A.** Head circumference measurements of 16p11.2 deletion (16p_del) without macrocephaly. **B.** Immunostaining for the NPC markers, Nestin and Pax6, in NPCs differentiated from 16p11.2 deletion without macrocephaly after 12 days of directed differentiation. Scale bar: 50 μm. **C.** mRNA expression levels for *pax6* in 16p_del lines without macrocephaly after 12 days of directed differentiation into NPCs compared to control lines. Data represent fold change relative to undifferentiated control human iPSCs. **D.** Quantitative RT-PCR of *cd47* mRNA expression in NPCs derived from 16p_del individuals without macrocephaly compared to control subjects. Data represent fold change relative to undifferentiated control human iPSCs. **E.** Quantification of CD47 mean fluorescence intensities (MFI) of CD47 protein levels in 16p_del NPCs without macrocephaly compared to control NPCs. **F.** Quantification of cell surface calreticulin (CRT) MFI in 16p_del NPCs without macrocephaly and control NPCs. n=2 biological replicates per cell line (control, n=3; 16p_del no macrocephaly, n=2 cell lines). All data are mean ± SE and were subjected to a two-tailed student’s *t*-test.

**Supplementary Figure 4. Directed differentiation of 16p11.2 CNV and control iPSC lines into oligodendrocyte progenitor cells**.

**A.** Flow cytometry analyses for the oligodendrocyte progenitor cell markers, NG2 and O4, in the 16p11.2_dup, 16p11.2_del, and control iPSC lines after 50 days of directed differentiation.

**Supplementary Figure 5. CD47 expression is upregulated in 16p11.2 deletion in O4+ fluorescence activated cell sorted (FACS) cells**.

**A.** Quantitative RT-PCR showing mRNA expression levels of *cd47* in O4+ FACS-sorted cells in the 16p_del lines compared to control and 16p_dup lines following 50 days of directed differentiation. **B.** Quantitative RT-PCR of mRNA expression levels of several genes associated with the oligodendrocyte lineage, including *olig2*, *nkx2.2*, *olig1* and *ng2* in O4+ FACS-sorted cells. **C.** Quantitative RT-PCR showing mRNA expression levels of the neuronal (*tuj1*), astrocytic (*gfap*), and the mature oligodendrogenic (myelin basic protein, *mbp*) markers in O4+ FACS-sorted cells. **D.** Quantitative RT-PCR of mRNA expression levels of genes associated with the spinal cord (*cdx1*, *cdx2*, *cdx4*, *hoxb9*, *hoxb13*) in the 16p_dup, 16p_del, and control O4+ FACS-sorted cells following 50 days of directed differentiation. All data represent fold change relative to undifferentiated control human iPSCs. n=3 biological replicates per cell line (16p_dup, n=2; control, n=3; 16p_del, n=4 cell lines). All data are mean ± SE; P values were determined by one-way ANOVA followed by *post-hoc* Tukey HSD Test.

**Supplementary Figure 6. CD47 expression is not altered in OPCs derived from 16p11.2 deletion lines without macrocephaly**.

**A.** Flow cytometry analyses for the oligodendrocyte progenitor cell markers, NG2 and O4, in the 16p11.2_del individuals without macrocephaly after 50 days of directed differentiation. **B.** Quantitative RT-PCR of *cd47* mRNA expression in OPCs derived from 16p_del individuals without macrocephaly compared to control subjects. Data represent fold change relative to undifferentiated control human iPSCs. **C.** Quantification of CD47 mean fluorescence intensities (MFI) of CD47 protein levels in 16p_del OPCs without macrocephaly compared to control OPCs. **D.** Quantification of cell surface calreticulin (CRT) MFI in 16p_del OPCs without macrocephaly and control OPCs. n=2 biological replicates per cell line (control, n=3; 16p_del no macrocephaly, n=2 cell lines). All data are mean ± SE and were subjected to a two-tailed student’s *t*-test.

**Supplementary Figure 7. Representative controls for CD47 and Calreticulin mean fluorescence intensity assessments**.

Representative histograms showing the mean fluorescence intensities (MFI) of CD47 in unstained **(A)** hiPSCs, **(B)** NPCs, and **(C)** OPCs. Representative histograms showing the MFI of cell surface calreticulin (CRT) in unstained **(D)** NPCs, and **(E)** OPCs.

**Supplementary Table 1. Clinical characteristics of 16p11.2 CNV subjects and controls.**

Summary of the clinical and phenotypic data of the 16p11.2 CNV subjects included in the study.

**Supplementary Table 2. Primer sequences for qRT-PCR.**

Primer sequences included in the study.

