## Supplementary figures and tables for "Overexpression of CD47 is associated with brain overgrowth in 16p11.2 deletion syndrome"

### Supplementary Figure 1

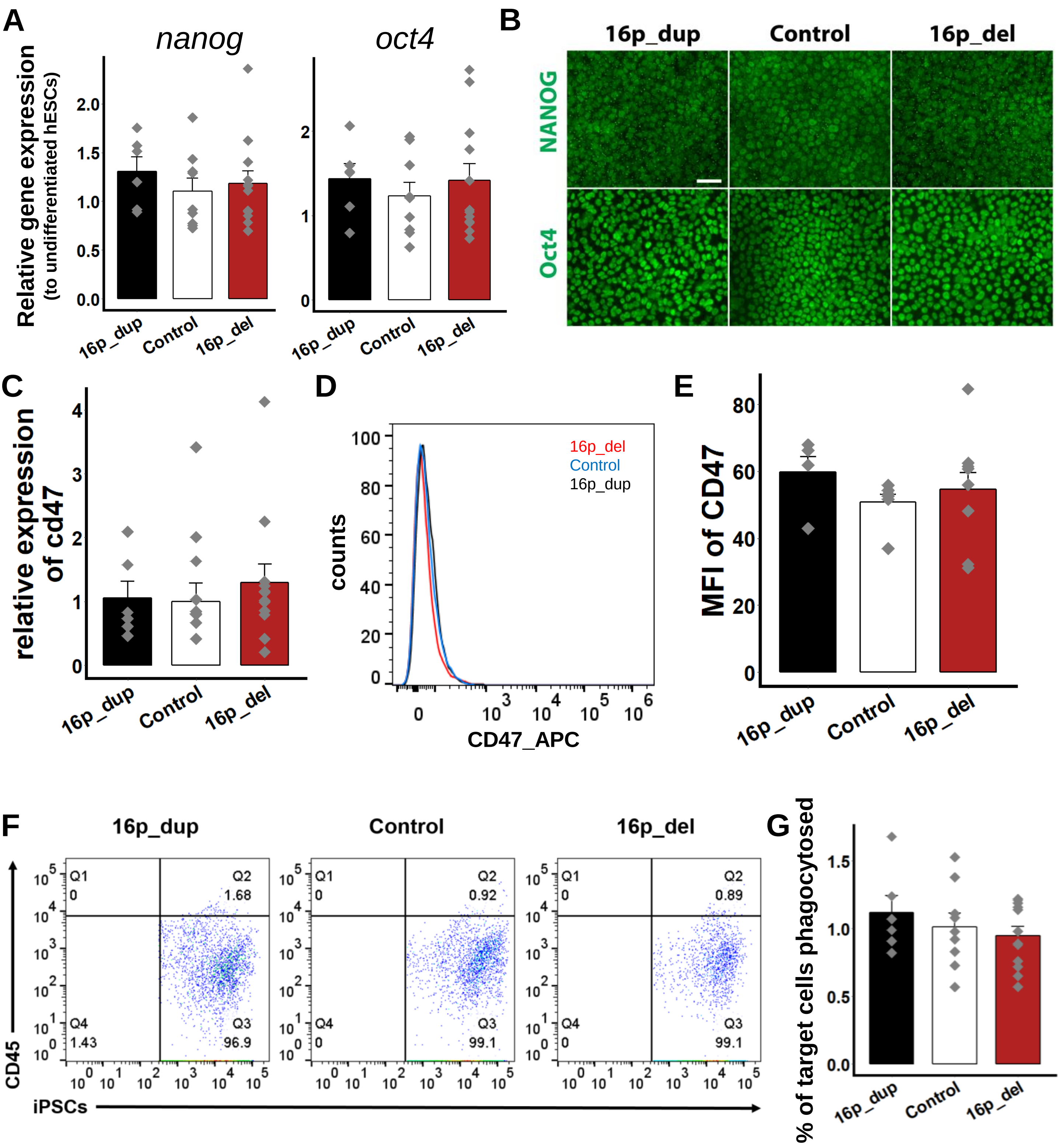

### Supplementary Figure 2

A

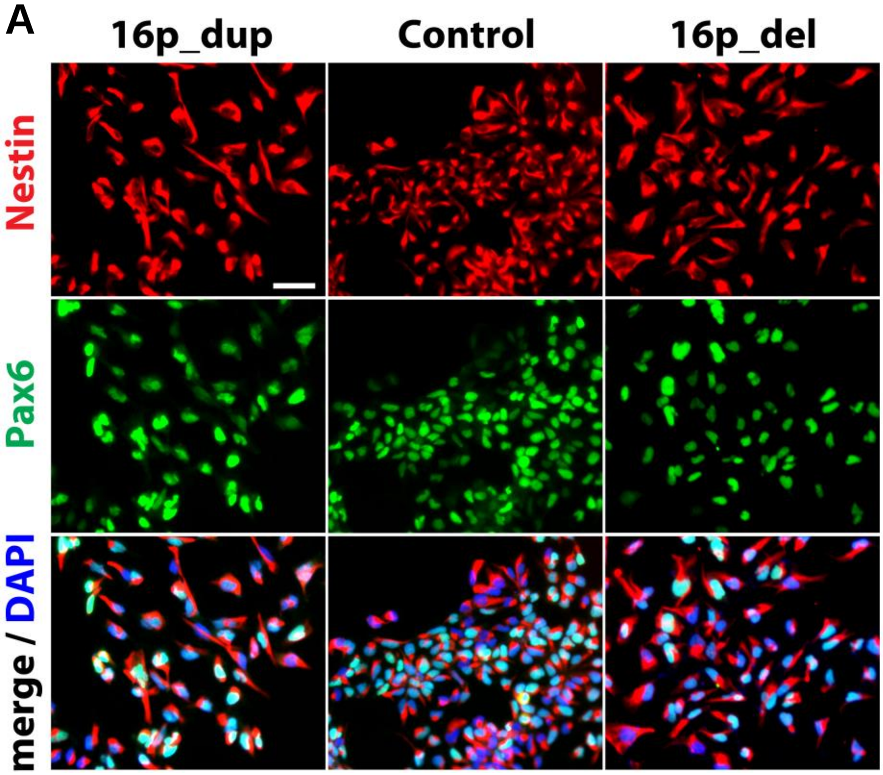

### Supplementary Figure 3

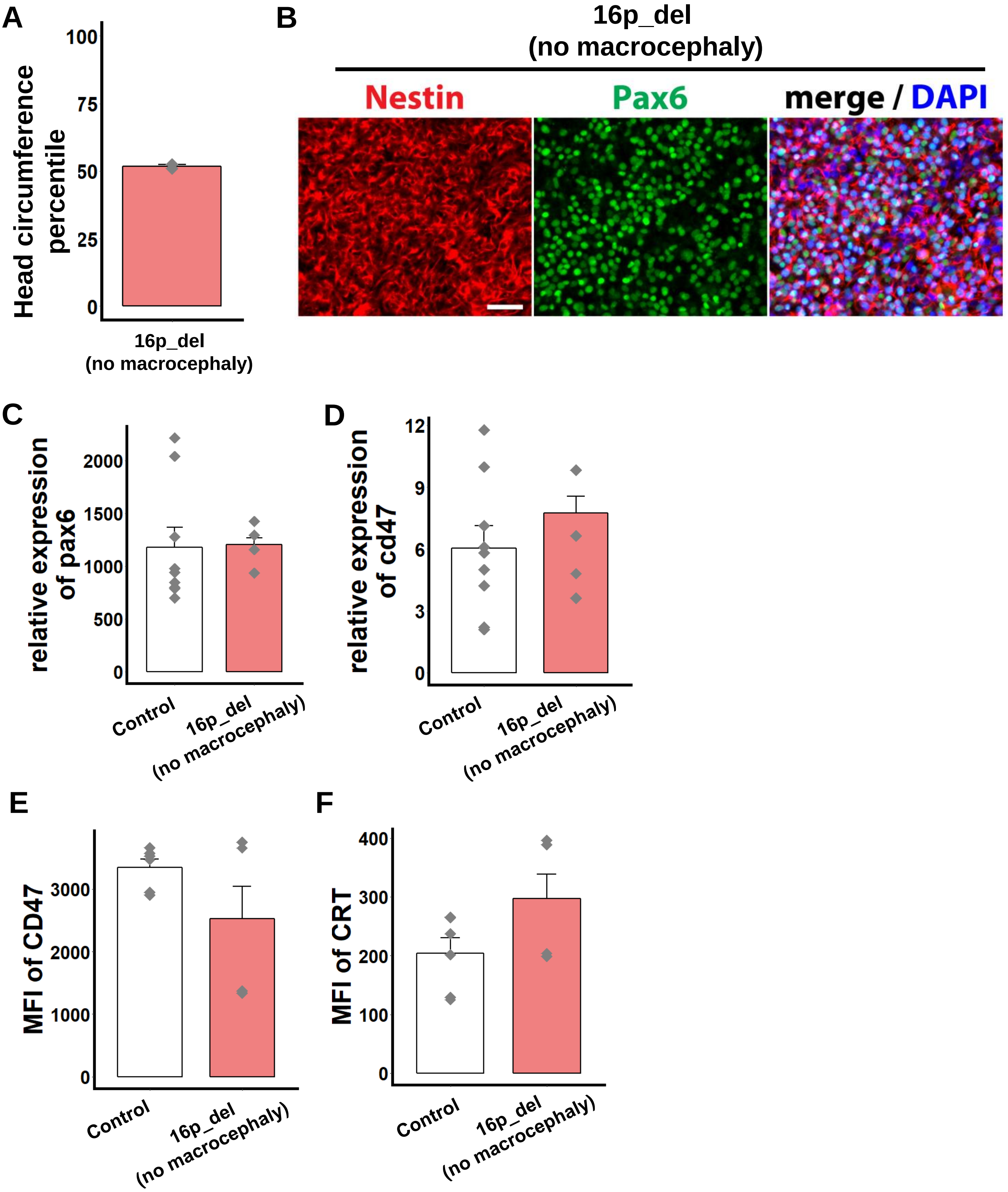

### Supplementary Figure 4

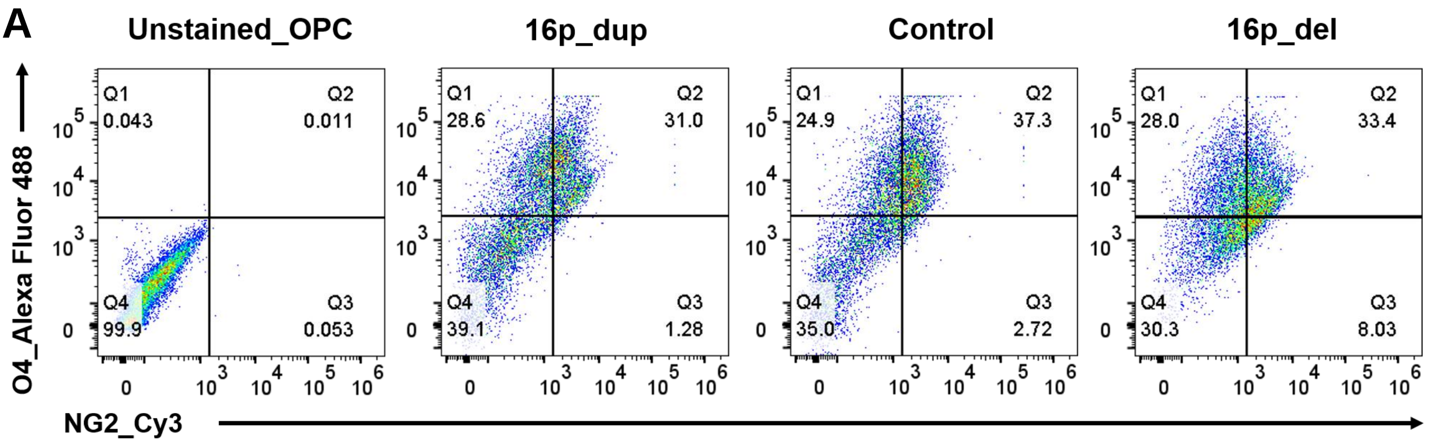

### Supplementary Figure 5

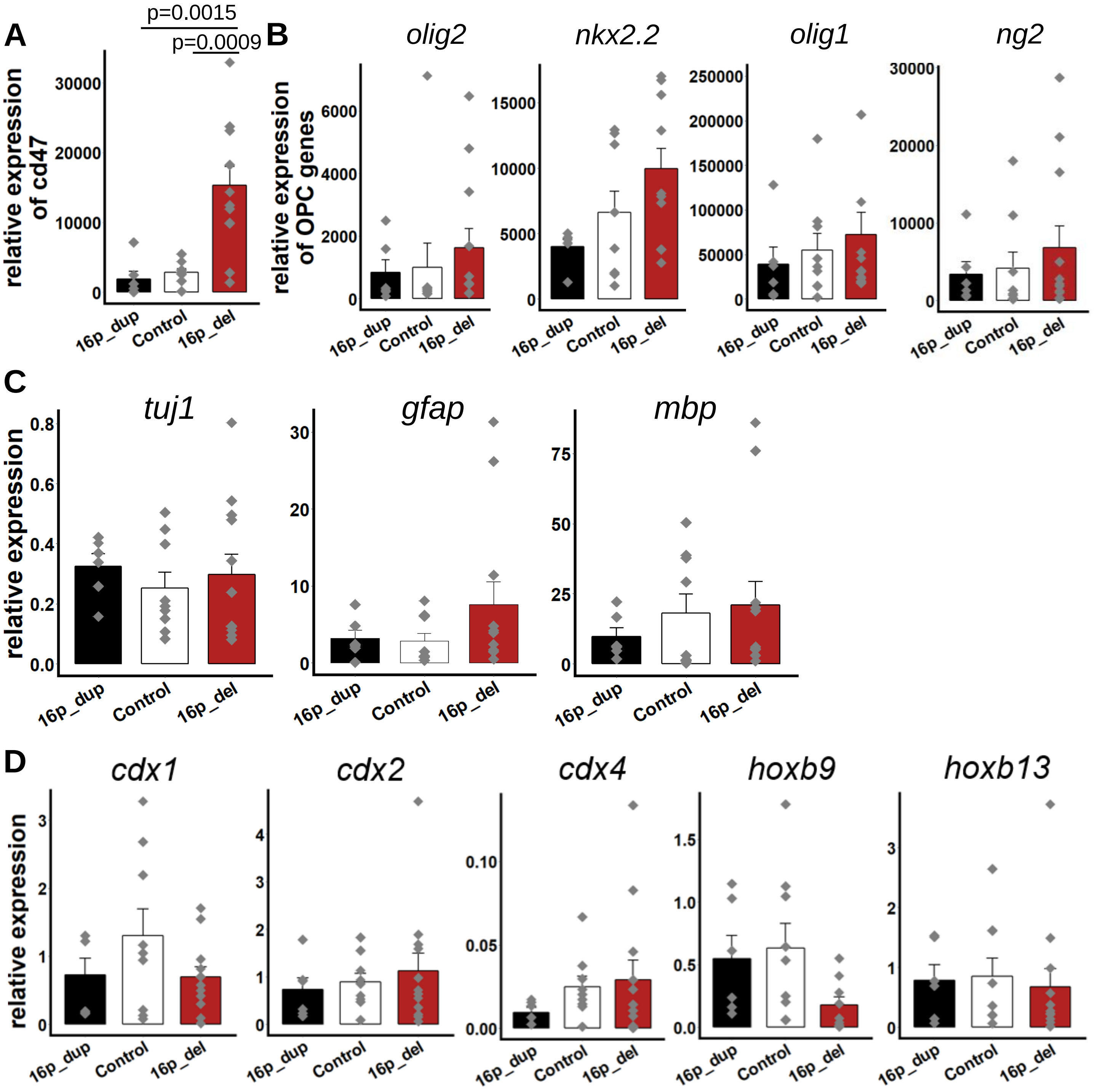

### Supplementary Figure 6

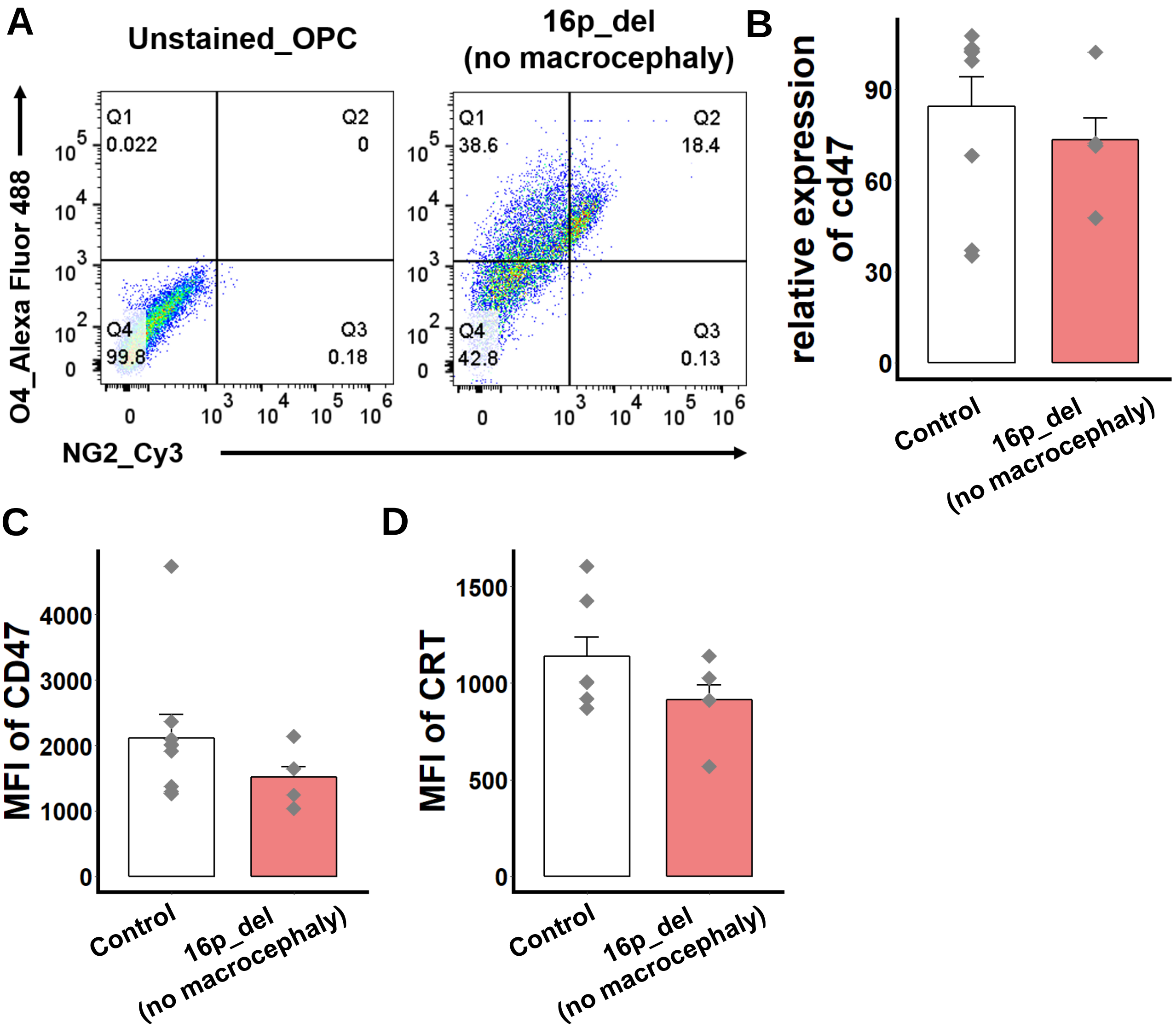

### Supplementary Figure 7

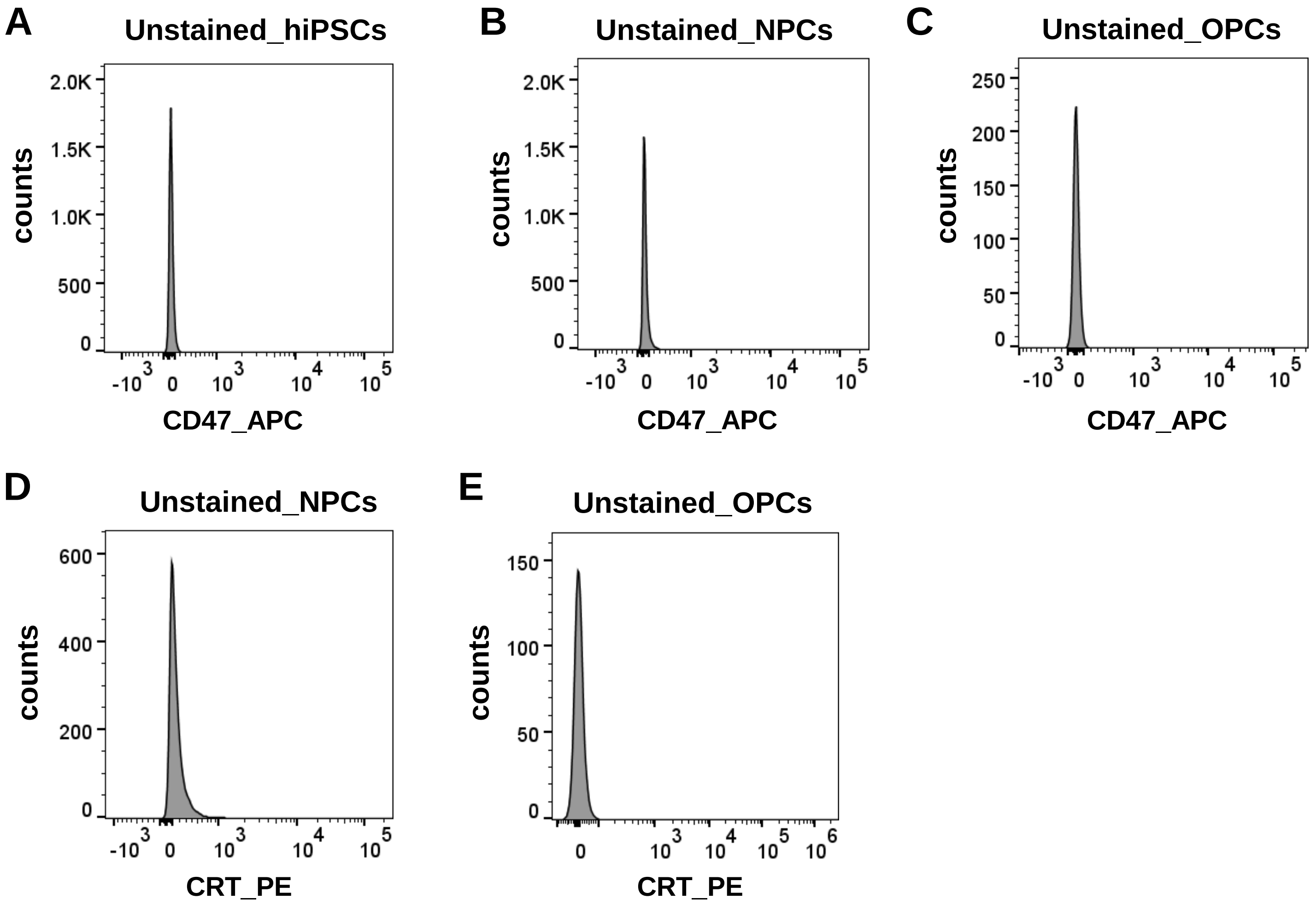

**Supplementary Table 1. Clinical characteristics of 16p11.2 CNV subjects and controls**

| Subject ID | CNV status | Inheritance | Diagnosis | Age of Evaluation (months) | Head Circumference Percentile |
|---|---|---|---|---|---|
| 14756.x16 | 16p11.2-duplication | inherited | Non-spectrum-diagnosis | 45 | 12 |
| 14756.x9 | 16p11.2-duplication | unknown | No diagnosis | 315 | 73 |
| 14739.x3 | 16p11.2-deletion | unknown | Intellectual disability | 95 | 99 |
| 14765.x2 | 16p11.2-deletion | unknown | Intellectual disability | 152 | 91 |
| 14799.x1 | 16p11.2-deletion | de-novo | Autism spectrum disorder | 174 | 99 |
| 14824.x13 | 16p11.2-deletion | de-novo | Autism spectrum disorder | 172 | 99 |
| 14763.x7 | 16p11.2-deletion | de-novo | Non-spectrum-diagnosis | 62 | 48 |
| 14746.x8 | 16p11.2-deletion | inherited | Autism spectrum disorder | 119 | 57 |
| 8343 | normal | - | No diagnosis | - | - |
| 2788.3 | normal | - | No diagnosis | - | - |
| 511 | normal | - | No diagnosis | - | - |

**Supplementary Table 2. Primer sequences for qRT-PCR.**

| Human Gene | Forward Sequence | Reverse Sequence |
|---|---|---|
| *gapdh* | GGAGCGAGATCCCTCCAAAAT | GGCTGTTGTCATACTTCTCATGG |
| *cdipt* | TTCTTTCTACTTCATGCCCTG | CCCAAACCGGGTTCCTT |
| *coro1a* | TGTCAACCCTAAGTTTGTGG | TTCTTGTCCACACGTCCA |
| *ppp4c* | GCACTGAGATCTTTGACTACC | TCGGTCGATTGTCCGAAT |
| *qprt* | CTGGTGCCGACCTTGT | GAACTGGGCCTTCAGCA |
| *ypel3* | CAGGCCTACTTGGATGATTG | TGAGTTGAAGAGGTAGGCA |
| *pax6* | GCAGATGCAAAAGTCCAGGTG | CAGGTTGCGAAGAACTCTGTTT |
| *cd47* | AGAATTCACGTTTTGTAATGACACT | GTGGGGACAGTGGACTTGTT |
| *olig2* | TGCGCAAGCTTTCCAAGAT | CAGCGAGTTGGTGAGCATGA |
| *nkx2.2* | GACAACTGGTGGCAGATTTCGCTT | AGCCACAAAGAAAGGAGTTGGACC |
| *olig1* | TGCTATGACCTTTCCGCAGT | ACACCGTCAGGAAACAAGGT |
| *ng2* | CTTCATCCACGATGGCTCTGA | TGGGCAGGAGGTATGTTTGG |
| *pdgfrα* | GGGCACGCTCTTTACTCCAT | ACAGCCTAAGACCAGGAACG |
| *mbp* | GGCAAGGTACCCTGGCTAA | GGGTGGTGTGAGTCCTTGTA |
| *gfap* | AAGATCCACGAGGAGGAGGTT | TGCGTGCGGATCTCTTTCAG |
| *tuj1* | GGCCAAGGGTCACTACACG | GCAGTCGCAGTTTTCACACTC |
| *cdx1* | GGTTACAGAATCACAGCCCTC | CTTGGTCCGAATAAAGTCCTC |
| *cdx2* | GGGCTCTCTGAGAGGCAGGT | CCTTTGCTCTGCGGTTCTG |
| *cdx4* | AGTCTGGGGCTCACCCTAC | CTGTGCCCATTGTACTAGACG |
| *hoxb9* | CCGTCTACCACCCTTACATCC | CGTAGCCGGGTCTTTGATTAG |
| *hoxb13* | AGCTCCCGTGCCTTATGGTTA | GGCTGGTAGGTTCCCGGATA |
